# Self-normalizing learning on biomedical ontologies using a deep Siamese neural network

**DOI:** 10.1101/2020.04.23.057117

**Authors:** Fatima Zohra Smaili, Xin Gao, Robert Hoehndorf

**Affiliations:** King Abdullah University of Science and Technology (KAUST), Computational Bioscience Research Center (CBRC), Computer, Electrical & Mathematical Sciences and Engineering (CEMSE) Division, Thuwal 23955, Saudi Arabia

## Abstract

**Motivation:** Ontologies are widely used in biomedicine for the annotation and standardization of data. One of the main roles of ontologies is to provide structured background knowledge within a domain as well as a set of labels, synonyms, and definitions for the classes within a domain. The two types of information provided by ontologies have been extensively exploited in natural language processing and machine learning applications. However, they are commonly used separately, and thus it is unknown if joining the two sources of information can further benefit data analysis tasks.

**Results:** We developed a novel method that applies named entity recognition and normalization methods on texts to connect the structured information in biomedical ontologies with the information contained in natural language. We apply this normalization both to literature and to the natural language information contained within ontologies themselves. The normalized ontologies and text are then used to generate embeddings, and relations between entities are predicted using a deep Siamese neural network model that takes these embeddings as input. We demonstrate that our novel embedding and prediction method using self-normalized biomedical ontologies significantly outperforms the state-of-the-art methods in embedding ontologies on two benchmark tasks: prediction of interactions between proteins and prediction of gene–disease associations. Our method also allows us to apply ontology-based annotations and axioms to the prediction of toxicological effects of chemicals where our method shows superior performance. Our method is generic and can be applied in scenarios where ontologies consisting of both structured information and natural language labels or synonyms are used.

**Availability:** https://github.com/bio-ontology-research-group/Ontology-based-normalization

## 1 Introduction

Biomedical ontologies have shown their usefulness in performing different types of analysis tasks and have contributed significantly to accelerating research in the biomedical field. Ontologies do not serve anymore exclusively as information models that formally represent the massive amounts of biomedical knowledge and its complex relations, but they have also shown potential in improving machine learning algorithms in several tasks including disease–gene prioritization (Schlicker *et al.*, 2010; Ortutay and Vihinen, 2008), protein function prediction (Kulmanov *et al.*, 2017), and protein-protein interaction prediction (De Bodt *et al.*, 2009; Smaili *et al.*, 2018a; Martin *et al.*, 2011), as well as in natural language processing (Muller *et al.*, 2004; Rzhetsky *et al.*, 2000), querying of biomedical data (Shah *et al.*, 2006), and similar.

Specifically, in biomedical applications, ontologies can contribute to improving machine learning based prediction tasks mainly in two ways. First, they can provide useful features for the biomedical entities of interest (e.g., protein functions using the gene ontology (GO) (Kulmanov *et al.*, 2017), disease phenotype associations based on the Human Phenotype Ontology (HPO) (Smaili *et al.*, 2018b)). Second, they can be used as domain-specific background knowledge that provide annotation property assertions which relate classes to their labels, synonyms, descriptions, etc. In both cases, the predictive performance of the machine learning model can significantly improve if the use of ontologies is combined with existing literature that can complement the information encoded in the ontology annotation assertions (Smaili *et al.*, 2018b).

It remains a challenge to combine ontologies and the information contained about the same or related phenomena in the literature so that the information contained in both can be exploited as background knowledge when building machine learning models. In particular, it is challenging to identify when a class in an ontology has been mentioned in literature and then build a model that can combine information about a class obtained from literature and from the knowledge contained within an ontology.

There are several methods available that use text mining methods to identify the mentions of ontology classes in text (Byrd and Ravin, 1999; Faure and Nédellec, 1998; Maedche and Staab, 2000; Morin, 1999; Krauthammer and Nenadic, 2004; Rebholz-Schuhmann *et al.*, 2007; Ma *et al.*, 2012; Rajpathak and Singh, 2013; Serra *et al.*, 2014). Here, we propose a method that annotates a literature corpus with classes in an ontology and then normalizes the corpus to the classes within the ontology by replacing each mention of an ontology class in text with the identifiers of the class. We apply this normalization method to biomedical literature as well as to the descriptions of classes within an ontology itself, thereby “self-normalizing” the natural language annotation assertions within an ontology.

We apply these normalization methods so that we can utilize literature and ontologies jointly in machine learning models. Ontology-based machine learning methods mainly use the formal axioms and entity-class annotations without considering natural language information (Smaili *et al.*, 2018a; Kulmanov *et al.*, 2019; Holter *et al.*, 2019). At the same time, there are a number of methods which apply learning on literature directly to perform biomedical analysis and prediction tasks (Kim *et al.*, 2019; Naeem *et al.*, 2010; Wong and Shatkay, 2013). We use our ontology-normalized dataset to develop a method that can jointly learn from literature and ontologies through; our method exploits the fact that ontology-based normalization makes the literature and ontologies overlap on the token-level. We use transfer learning and language models to develop a joint learning method that can generate embeddings for classes in ontologies and entities that are characterized through ontologies. Using a novel deep Siamese neural network architecture, we significantly improve prediction of relations between biological entities based on the embeddings we generate. The results from our experiments show that the combination of ontologies with information in literature and their own annotations, in combination with a deep Siamese neural network, can efficiently characterize semantic similarity and improve the performance in several prediction tasks, including prediction of interactions between proteins, prioritizing gene–disease associations, and prediction the toxicological effects of chemicals.

## 2 Materials and methods

### 2.1 Ontologies

We downloaded the Gene Ontology (GO) (Ashburner *et al.*, 2000) in Web Ontology Language (OWL) (Grau *et al.*, 2008) format from http://www.geneontology.org/ontology/ on April 14, 2018. This version of GO contains 107,762 logical axioms. We also downloaded the GO protein annotations from the UniProt-GOA website (http://www.ebi.ac.uk/GOA) on Dec 2, 2018. All associations with evidence code `IEA` were filtered, which resulted in a total of 3,474,539 associations for 749,938 unique proteins.

GO-Plus (downloaded from http://purl.obolibrary.org/obo/go/extensions/go-plus.owl) is an extension of GO that contains, in addition to all the logical axioms of GO, inter-ontology axioms that link GO classes to other external biomedical ontologies, in particular: ChEBI (The Chemical Entities of Biological Interest ontology) (Degtyarenko *et al.*, 2007), PO (The Plant Ontology) (Jaiswal *et al.*, 2005), CL (The Cell Ontology) (Bard *et al.*, 2005), PATO (Phenotype and Trait Ontology) (Gkoutos *et al.*, 2005, 2017), the Uberon ontology (Mungall *et al.*, 2012), SO (The Sequence Ontology) (Eilbeck *et al.*, 2005), FAO (Fungal gross anatomy), OBA (Ontology of Biological Attributes), NCBITaxon (NCBI organismal classification), CARO (Common Anatomy Reference Ontology) (Haendel *et al.*, 2008) and PR (Protein Ontology) (Natale *et al.*, 2010).

We downloaded the PhenomeNET ontology (Hoehndorf *et al.*, 2011; Rodríguez-García *et al.*, 2017) in the OWL format from the AberOWL repository http://aber-owl.net (Hoehndorf *et al.*, 2015) on February 21, 2018. PhenomeNET is a cross-species phenotype ontology that combines phenotype ontologies, anatomy ontologies, GO, and several other ontologies in a formal manner (Hoehndorf *et al.*, 2011).

We downloaded the Human Phenotype (HPO) ontology in the OWL format from http://purl.obolibrary.org/obo/hp.owl on February 7, 2019. This ontology contains 34,373 unique classes and 74,426 logical axioms. The HPO ontology formally describes the phenotypic abnormalities frequently encountered in human monogenic diseases (Robinson *et al.*, 2008).

We downloaded the Mammalian Phenotype (MP) ontology in the OWL format from http://purl.obolibrary.org/obo/mp.owl on February 7, 2019. The MP ontology contains 33,356 unique classes and 74,119 logical axioms. The MP ontology classifies and describes phenotypic information related to mice and other mammalian species that are used as models to study human diseases (Smith *et al.*, 2005; Smith and Eppig, 2009).

### 2.2 Normalization Methods

#### 2.2.1 Ontology-based dictionaries

To perform the ontology-based normalization of both the PMC articles and the ontology, we extract the labels and synonyms from the annotation property assertions of the ontology to create a lexical dictionary that maps a sequence of words to an ontology class IRI while filtering the most common words or sequence of words based on their frequency in the British National Corpus (http://www.natcorp.ox.ac.uk/). We have created a dictionary based on GO-plus for PPI prediction and a dictionary based on MP, and HPO ontologies for the gene–disease association prediction and the chemical–disease association prediction. For each ontology, the dictionary maps each ontology class (IRI) to the sequence of words that can refer to it, which have been extracted from the labels and synonyms (`rdfs:label`) and synonyms (`hasExactSynonym, hasRelatedSynonym, hasBroadSynonym, hasNarrowSynonym`) of each class based on the information available in the ontology annotation property assertions. To allow for morphological variations, we convert all words to their lower case while ignoring white spaces, hyphens and punctuation. If a class is mapped to only one word which happens to be in the list of the most common words in English according to the British National Corpus, this mapping is ignored. The created dictionaries are available at: https://github.com/bio-ontology-research-group/Ontology-based-normalization.

#### 2.2.2 Whatizit

The lexical dictionaries created are then used as inputs to Whatizit (Rebholz-Schuhmann *et al.*, 2007) to annotate PMC articles using biomedical ontology classes. Whatizit is a named entity recognition tool used to annotate text corpora by identifying biomedical classes and terminologies obtained either from known biomedical databases (e.g. Swiss-Prot, DrugBank, etc) or from dictionaries provided by the user (Rebholz-Schuhmann *et al.*, 2007). The strength of Whatizit lies in its ability to annotate large files of text relatively fast and in allowing for morphological variations during the normalization. We use Whatizit in this work to annotate all open-access full-text PMC articles obtained from ftp://ftp.ncbi.nlm.nih.gov/pub/pmc on January 10, 2020 using the ontology-based dictionaries. The Whatizit annotation is based on dictionaries that map genes and disease IDs to their labels in order to annotate PMC articles.

#### 2.2.3 Self-normalization

To make better use of the natural language information encoded in the ontology itself, we annotate the natural language definitions of classes with the ontology classes to detect further relations between biomedical classes. In particular, we annotate all values of class definition annotation property assertions in the ontology by using the created ontology-based dictionaries to directly replace all the existing occurrences of labels and synonyms in any description axiom with their corresponding ontology class IDs.

### 2.3 Embedding methods

#### 2.3.1 Text-Based embeddings

To produce vector representations of biomedical entities based on literature only as a baseline method, we use Whatizit to annotate PMC articles with protein Uniprot IDs for PPI prediction, MGI gene IDs and OMIM disease IDs for gene–disease association predictions, and CTD chemical IDs and OMIM disease IDs for chemical–disease associations. We then apply the skip-gram model from word2vec to produce the vector representations of the biological entities based on literature only without including ontology-based information. The obtained vectors are then given as inputs to the neural network in order to predict PPI or gene–disease associations.

#### 2.3.2 OPA2Vec

OPA2Vec (Smaili *et al.*, 2018b) is a method that uses ontologies to produce vector embeddings of ontology classes and the entities they annotate. OPA2Vec uses logical axioms as well as annotation property axioms from the ontology, in addition to the known ontology-based associations of biological entities (e.g. protein-GO associations). These annotation axioms use natural language to describe different properties of the ontology classes (labels, descriptions, synonyms, etc.) and they, therefore, form a rich corpus of text for Word2vec. To provide the Word2Vec model with background knowledge on the ontology classes described by the annotation properties, OPA2Vec pre-trains the model on a corpus of biomedical text. Entity-class annotations are also used as an additional source of information to produce the ontology-based embeddings of biological entities.

### 2.4 Siamese neural network

To predict associations between the embeddings of a pair of biological entities, we use a deep Siamese neural network. Siamese neural networks were first proposed in 1994 as an algorithm for signature verification (Bromley *et al.*, 1994). A Siamese neural network consists of a pair of twin networks in parallel, with similar parameter values, which take distinct samples and are joined by a similarity optimization function (Koch *et al.*, 2015). The Siamese neural network we use in this paper consists of two twin deep feed-forward multi-layer perceptron (MLP) networks with three hidden layers. The MLPs are joined by a contrastive loss function that computes the similarity between the highest-level feature representation of each entity produced by each MLP side. This is accomplished by taking the dot product of the feature representations produced by the twin MLPs followed by a top fully connected layer of 200 neurons and a binary output layer. The parameters of the Siamese neural network are trained using the cross entropy loss and back propagation. Figure 1 shows the architecture of the neural network used in this work. To alleviate the issue of over-fitting, we use a dropout layer in each MLP as well as an early stopping strategy. To reduce data bias, we over-sample the least frequent diseases in the training data.

**Fig. 1:**
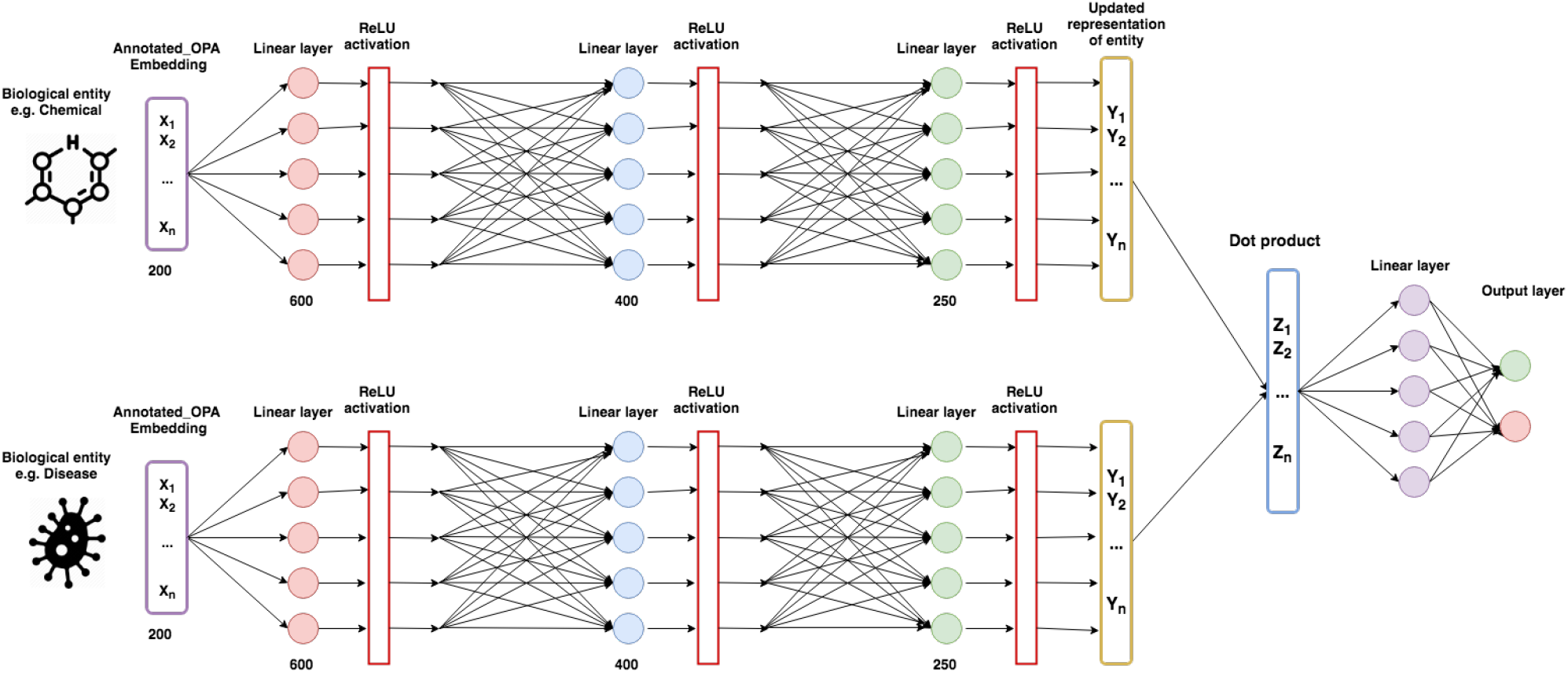
Architecture of the deep Siamese neural network, we use to predict associations between the embeddings of two biological entities. The Siamese neural network consists of two identical MLP networks that share the same architecture, the same parameters and the same weights.

### 2.5 Datasets

#### 2.5.1 Functional protein interactions

To evaluate our work, we predict functional protein interactions on three different organisms: human, yeast (*S. cerevisiae*), and *Arabidopsis thaliana*. The datasets for all three organisms were obtained from the STRING database (Szklarczyk *et al.*, 2017)(http://string-db.org). We filtered out interactions with score less than 700 to only keep interactions with strong evidence. The human dataset contains 19,577 proteins and 11,353,057 interactions; the yeast dataset contains 6,392 proteins and 2,007,135 interactions; and the Arabidopsis dataset contains 10,282,070 interactions for 13,261 proteins.

#### 2.5.2 Gene–disease associations

To further evaluate our method, we predict gene–disease associations. The first dataset used in this experiment is the mouse phenotype annotations obtained from the Mouse Genome Informatics (MGI) database (Smith and Eppig, 2015) on February 21, 2018 with a total of 302,013 unique mouse phenotype annotations. The second dataset used for this experiment is the disease to human phenotype annotations obtained on February 21, 2018 from the Human Phenotype Ontology (HPO) database https://hpo.jax.org/ (Robinson *et al.*, 2008). Our analysis is limited to the OMIM diseases only which results in a total of 78,208 unique disease-phenotype associations. To validate our prediction, we use the `MGI_DO.rpt` file from the MGI database to obtain 9,506 mouse gene-OMIM disease associations and 13,854 human gene-OMIM disease associations. To map mouse genes to human genes we use the `HMD_HumanPhenotype.rpt` file from the MGI database.

#### 2.5.3 Chemical–disease association

To predict associations between chemicals and diseases, we use the Comparative Toxicogenomics Database (CTD)(http://ctdbase.org) (Davis *et al.*, 2016) to obtain 108,783 unique chemical–disease associations between 7,248 unique diseases and 10,572 unique chemicals, downloaded on May 5, 2019. Among these chemical–disease associations, 34,573 are therapeutic associations while 62,915 are toxic (marker) associations. To create a balanced dataset, we subsample the negative pairs from all the remaining chemical–disease pairs that are not included in the CTD dataset. To create our ontology association file, we use GO functional annotations of chemicals and phenotypes annotations for diseases. We obtain a total of 5,416,206 chemical–GO associations from the CTD library, 78,208 disease–phenotype associations from the Human Phenotype Ontology (HPO) database (Robinson *et al.*, 2008), and 124,214 disease–phenotype associations from the disease ontology (Schriml *et al.*, 2011).

## 3 Results

### 3.1 Ontology-based normalization of natural language

Embedding the classes, relations, and instances in ontologies can provide useful features for predictive models that rely on background knowledge, and these embeddings can incorporate ontology axioms as well as natural language annotations such as labels and definitions (Kulmanov *et al.*, 2019; Liu-Wei *et al.*, 2019; Althubaiti *et al.*, 2019; Smaili *et al.*, 2018a). However, using the natural language information in ontologies can also add noise, in particular when labels or descriptions use complex terms, such as chemical formulas, which are not easy to recognize in natural language text (Smaili *et al.*, 2019).

We propose a novel method that more closely integrates ontologies and natural language text, including both literature and the labels, definitions, or synonyms contained within ontologies themselves. We first normalize the natural language text to the ontology using named entity recognition and normalization methods. This recognition and normalization step aims to detect and then “normalize” terms used to refer to a class in an ontology; here, normalization is the process of ensuring that the symbols used to refer to a class in an ontology in text or within an ontology are identical. To normalize natural language texts, either literature or the descriptions associated with classes in an ontology, we replace every occurrence of a term which refers to an ontology class with the identifier of that class in the ontology (i.e., the class IRI). To recognize mentions of ontology classes in text, we create a dictionary based on the labels and synonyms of the classes in the ontology, and use a dictionary-based named entity recognition and normalization method (Rebholz-Schuhmann *et al.*, 2007) to replace each mention of an ontology class with the class IRI. We apply this normalization method to any text used as background knowledge. An example is shown in Figure 2.

**Fig. 2:**
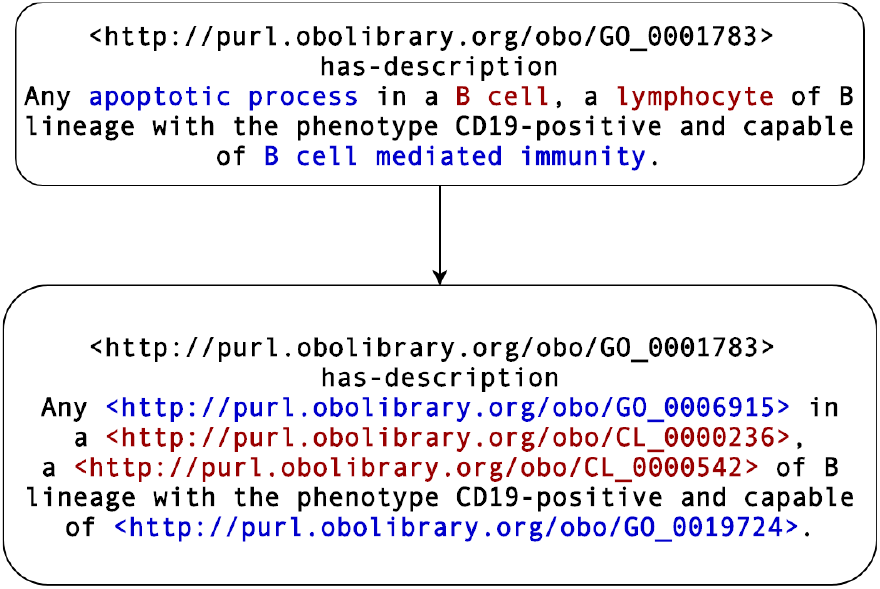
An example of the normalization within a class description in the GO ontology. In blue, we show examples of self-normalization where we replace the labels of GO classes (*apoptotic process* and *B cell mediated immunity*) with their corresponding IRI. In red, we show examples of GO normalization with the Cell Ontology (CL). The labels of CL classes (*B cell* and *lymphocyte*) are replaced with their IRI.

To learn vector representations of classes and entities, using only the information within ontologies, we modify the OPA2Vec method (Smaili *et al.*, 2018b). OPA2Vec uses a language model applied to a corpus consisting of asserted and deductively inferred axioms, as well as the annotation property assertions, within one ontology; we extend OPA2Vec with our ontology-based normalization method so that all string values used in an annotation property assertion are normalized to the class identifiers within the same ontology.

OPA2Vec has the ability to use transfer learning and thereby incorporate information from literature by pre-training a language model on a literature corpus and then training it further on the corpus generated from the axioms and annotation assertions of the ontology. We apply the same pre-training step in our method but normalize the literature corpus to the ontology before pre-training; we then apply the same transfer learning method as in OPA2Vec.

We expect the normalization of the literature and of ontology class descriptions to improve our ability of the embedding methods to capture previously undetected relations between ontology classes and, therefore, improve their performance in predicting associations between different biomedical entities. This prediction is based on detecting certain similarities within the embedding space, i.e., given two embeddings (such as the embeddings for two proteins, or for a gene and a disease), detect how similar they are (with respect to a certain similarity measure which is generated through supervised learning). We use a deep Siamese neural network architecture (Bromley *et al.*, 1994) that applies two multi-layer perceptron neural networks with shared weights on the two embeddings in a parallel fashion, followed by a dot product, to determine how similar these embeddings are. We use the output of this network as the prediction score for a similarity, or the existence of a relation, between the entities whose embeddings are used as input. The model is trained using a cross-entropy loss. Figure 1 provides an overview of the prediction model.

### 3.2 Protein interaction prediction

To determine whether our novel method can improve the performance of relation prediction, we apply it to a benchmark dataset for predicting interactions between proteins. Predicting interactions between proteins is an established benchmark for testing and comparing similarity-based methods, in particular methods that rely on annotations of proteins with ontologies (Pesquita *et al.*, 2008; Smaili *et al.*, 2018a,b) We use the GO-plus ontology (Consortium, 2014, 2016) classes to annotate the literature and its own annotation property assertions (i.e., the natural language definitions of classes), and use the rest of our workflow to produce embeddings of proteins based on GO and the protein-to-GO associations. The embeddings we generate are then used to predict interactions between proteins for human, yeast and *Arabidopsis Thaliana*, based on the prediction output of our Siamese neural network prediction model. The results from this experiment are referred to as *Annotated*_*OPA*2*Vec* in Figure 3 and Table 1. We also evaluate the impact of first normalizing the literature to the ontology (*Annotated*_*PMC*) and the class descriptions within the ontology annotation properties (*Annotated*_*metadata*) separately to assess the contribution of each component in improving the predictive performance of GO-plus for protein interactions. As a baseline method to which we compare our results, we produce vector representations of proteins from PMC articles using Word2vec without including any ontology information (referred to as *PMC*_*only* in the results). We also compare the obtained results to the vector representations obtained from GO-plus using the original OPA2Vec pipeline without any annotation property assertions or text normalization (referred to as *OPA*2*V ec*).

**Fig. 3:**
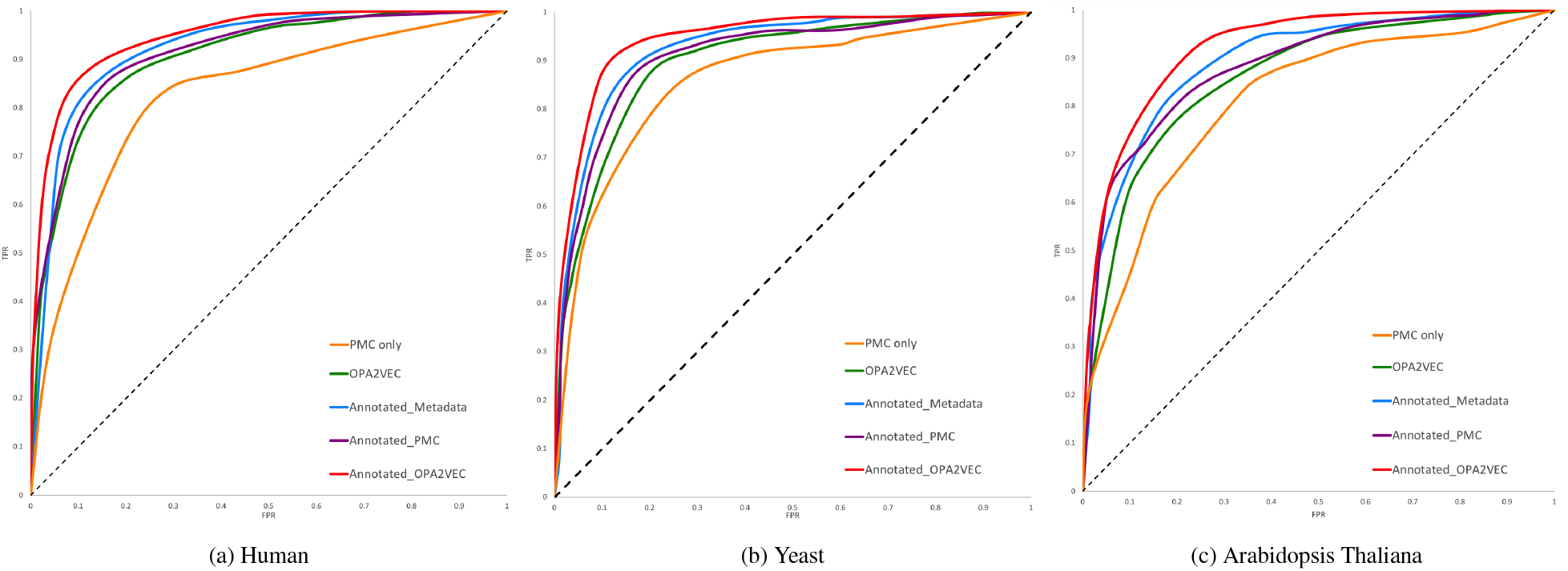
ROC curves for PPI prediction to show the contribution of the ontology-based annotation for literature and ontology descriptions.

**Table 1.** AUC values of ROC curves for PPI prediction to show the contribution of the ontology-based annotation for literature and ontology descriptions.

|  | Human | Yeast | Arabidopsis |
| --- | --- | --- | --- |
| *PMC_only* | 0.8171 | 0.8529 | 0.8089 |
| *OPA2Vec* | 0.8993 | 0.8951 | 0.8582 |
| *Annotated_Metadata* | 0.9187 | 0.9144 | 0.8726 |
| *Annotated_PMC* | 0.9093 | 0.9065 | 0.8650 |
| *Annotated_OPA2Vec* | **0.9384** | **0.9256** | **0.8882** |

Our results show that the annotation of both literature and the class descriptions in the ontology class descriptions performs best among all tested methods. Also, the performance improvement provided by the annotation properties is more significant than that of the literature annotation alone.

### 3.3 Gene–disease association prediction

As a second experiment to evaluate our novel method, we extend our analysis to the task of predicting gene–disease associations based on phenotype similarity using the PhenomeNet ontology. (Hoehndorf *et al.*, 2011; Rodríguez-García *et al.*, 2017) We use the annotations of human diseases with classes from the Human Phenotype Ontology (HPO) (Robinson *et al.*, 2008), and mouse genes using classes from the Mammalian Phenotype Ontology (MP) (Smith *et al.*, 2005). We then follow our prediction workflow and evaluate the performance of text and annotation property normalization using the HPO and MP ontology classes in predicting gene–disease associations for human and mouse, respectively. The results obtained are reported in Figure 4 and Table 2.

**Fig. 4:**
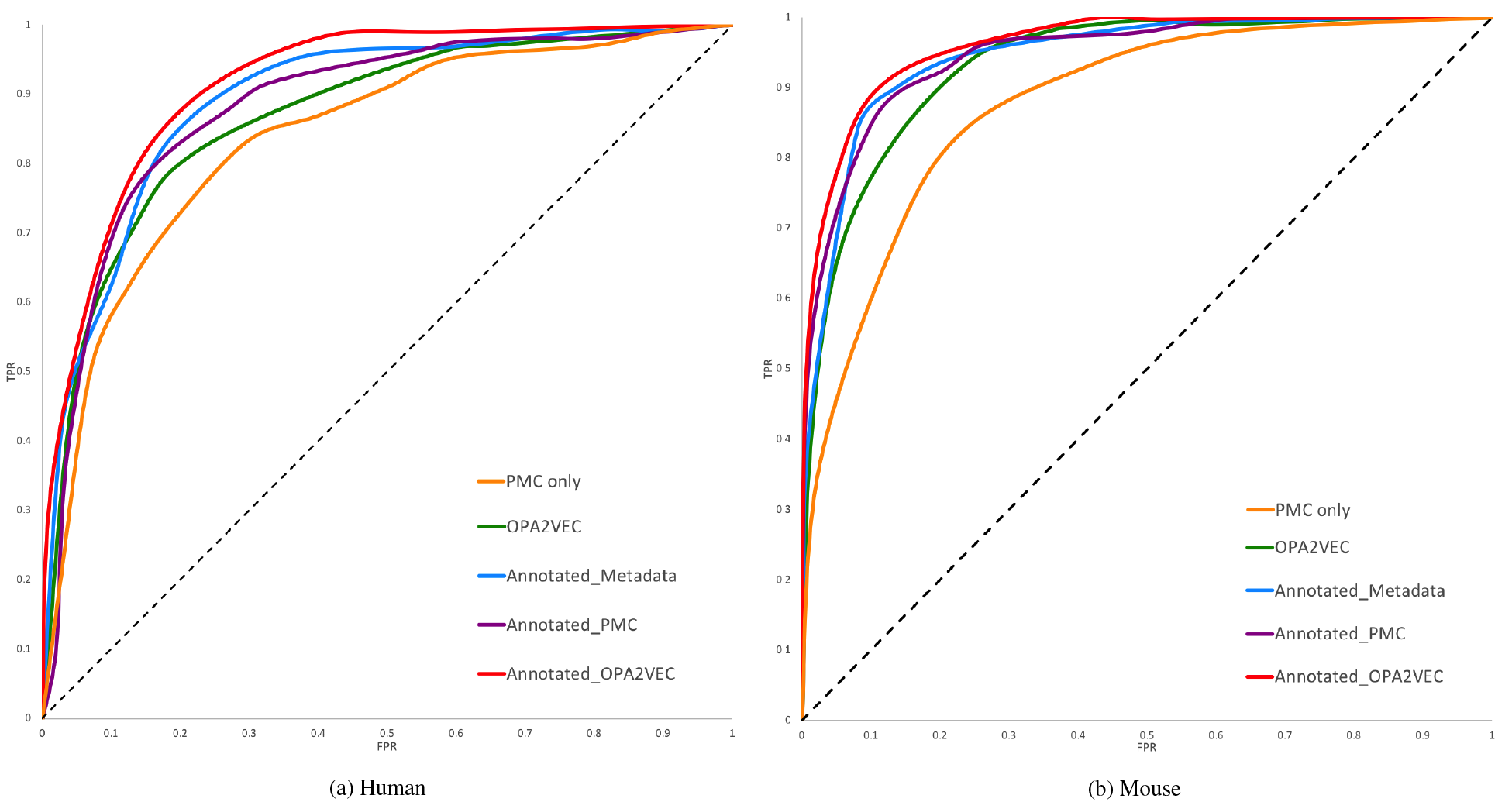
ROC curves for gene–disease association prediction.

**Table 2.**
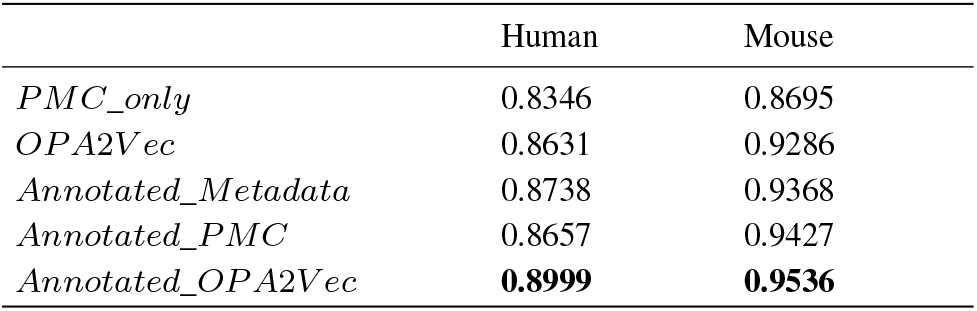
ROC curves for gene–disease association prediction to show the contribution of the ontology-based annotation for literature and ontology descriptions on human and mouse.

Similar to our evaluation of predicting interactions between proteins, the results from the gene–disease association prediction experiments show that the annotation of the class descriptions of the PhenomeNet ontology and the PMC articles with MP and HPO classes improves the prediction performance for both human and mouse. As our method makes use of information contained in literature, we directly compare our predictive performance to that of BeFree, a text-mining method that predicts gene–disease associations using biomedical named entity recognition (Bravo *et al.*, 2014, 2015). We compare our method with BeFree on the intersection of the Befree dataset and our gene–disease dataset consisting of 1,200 human diseases as reported in Table 3. Our results show that our normalization based method (Annotated_OPA2Vec) significantly outperforms BeFree (p-value of 0.031; Mann-Whitney U test) Figure 5 shows the overlapping gene–disease associations between BeFree, Annotated_OPA2Vec, OPA2Vec and the MGI database (ground truth). We find that the Annotated_OPA2Vec shares the highest numbers of positive associations with the MGI database.

**Fig. 5:**
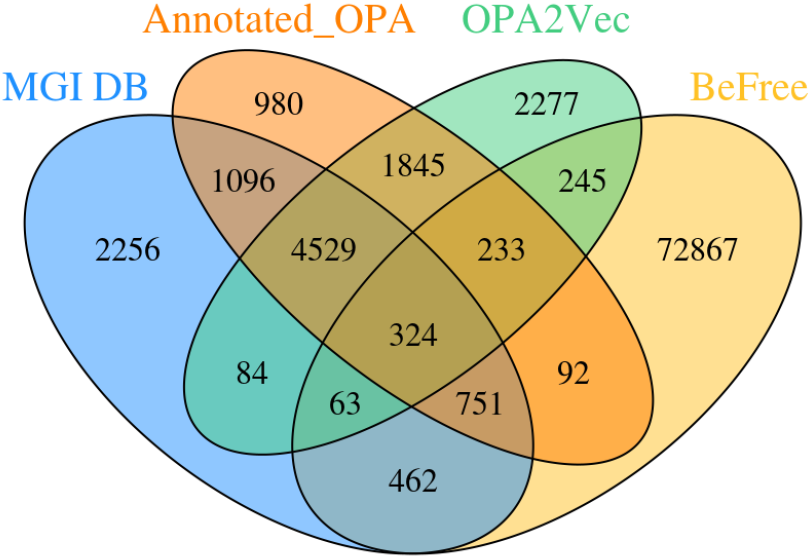
Overlapping of positive gene–disease associations between BeFree, Annotated_OPA2Vec, OPA2Vec and the MGI database (ground truth).

**Table 3.** AUC values of gene–disease association prediction on the intersection of our dataset and the BeFree dataset comparing the performance of BeFree to our methods.

|  | AUC |
| --- | --- |
| *BeFree* | 0.7550 |
| *PMC_only* | 0.7258 |
| *Annotated_Metadata* | 0.8717 |
| *Annotated_PMC* | 0.8543 |
| *Annotated_OPA2Vec* | **0.9071** |

As part of our validation, we predict gene–disease associations for orphan diseases, i.e., diseases which are suspected or known to have a genetic basis but where no gene association is currently known. Supplementary Table 1 shows predictions for those diseases for which an associated region is known and the gene predicted by us falls precisely within this region.

### 3.4 Chemical–disease association prediction

Related methods that combine ontology axioms and natural language information in a single machine learning model did not perform well when using information about chemicals, presumably due to the use of chemical formulas in class labels and descriptions (Smaili *et al.*, 2019). Through the use of ontology-based normalization, we hypothesize that we can somewhat overcome this limitation. One task for which rich information about relevant entities is available through ontologies is the identification of toxicological effects of chemicals, as contained in the Comparative Toxicogenomics Databases (CTD) (Davis *et al.*, 2016).

We represent chemicals through their GO functions and diseases through their phenotypes to generate embeddings, and we train our model using the information in CTD about chemical–disease associations. We compare the results of using PMC articles only without any ontology-based features, and the use of ontology-based associations with and without text normalization. The results from these experiments are shown in Figure 6 and Table 4. The results show that the literature and ontology normalization improves our ability to predict chemical–disease associations.

**Fig. 6:**
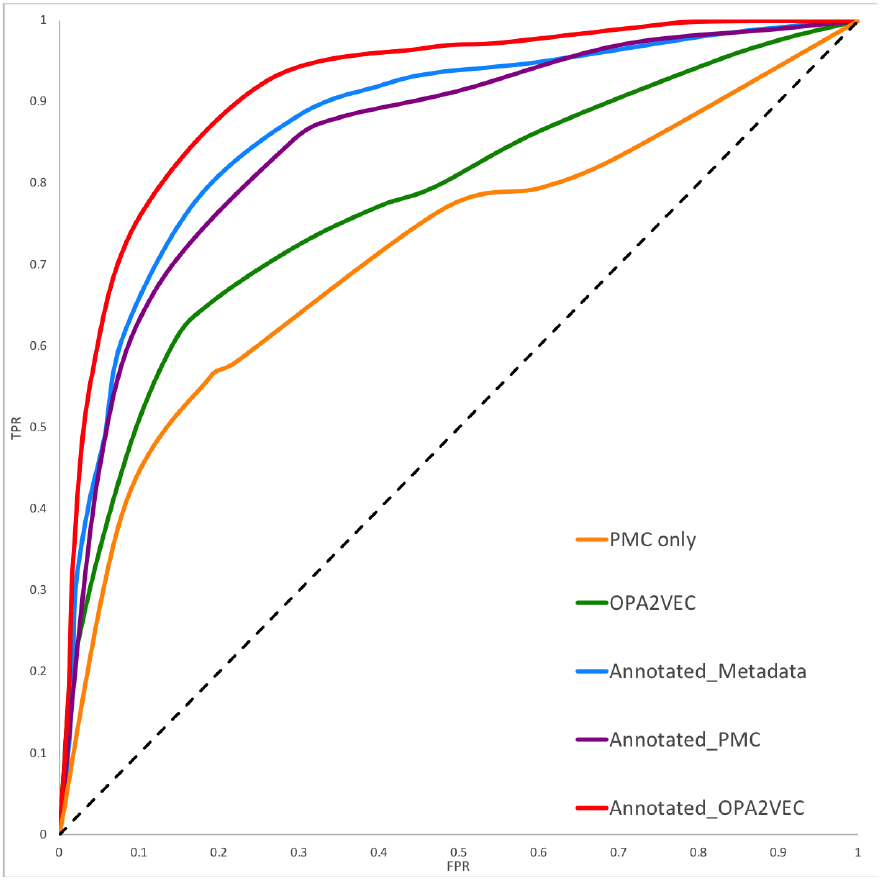
ROC curves for chemical–disease association prediction to show the contribution of the ontology-based annotation for literature and ontology descriptions based on the CTD dataset.

**Table 4.** AUC values of ROC curves for chemical–disease association prediction to show the contribution of the ontology-based annotation for literature and ontology descriptions based on the CTD dataset.

|  | AUC |
| --- | --- |
| *PMC_only* | 0.7114 |
| *OPA2Vec* | 0.7741 |
| *Annotated_Metadata* | 0.8694 |
| *Annotated_PMC* | 0.8501 |
| *Annotated_OPA2Vec* | **0.9104** |

To test whether our method is able to improve over the-state-of-the-art in predicting toxicological effects of chemicals, we compare our best-performing method, Annotated_OPA2Vec, in predicting chemical–disease associations to DigChem, a recent deep learning based method that predicts toxicological effects (chemical–disease associations) based on literature (Kim *et al.*, 2019). DigChem predictions do not report a confidence score, and therefore our performance comparison is based on accuracy, precision and recall values as reported in Table 5. The prediction results show that our normalization based method, Annotated_OPA2Vec, significantly outperforms DigChem (p-value of 0.046; Mann-Whitney U test) Figure 7 shows the overlapping chemical–disease associations between DigChem, Annotated_OPA2Vec, OPA2Vec and the CTD database (ground truth). We find that our normalization based method, Annotated_OPA2Vec, shares the highest number of associations with the curated CTD database (ground truth).

**Fig. 7:**
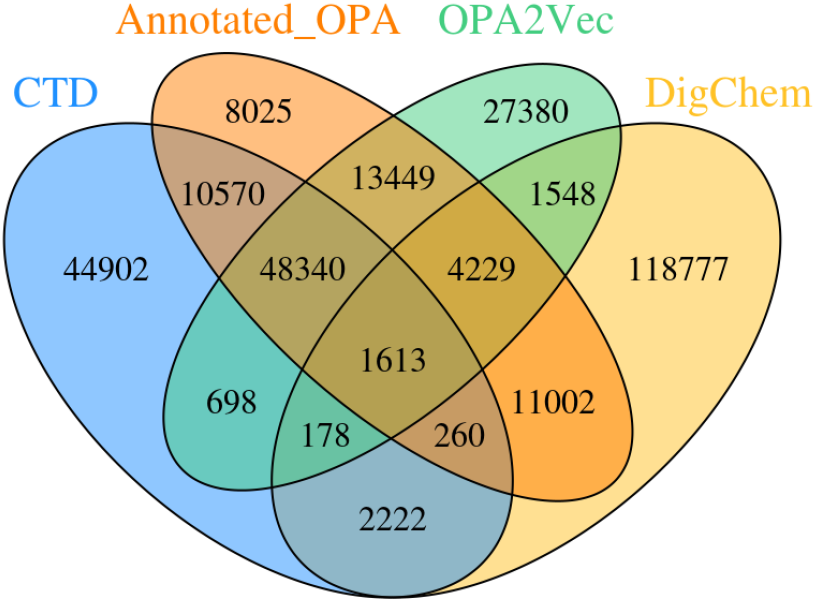
Overlapping positive chemical–disease associations between DigChem, Annotated_OPA2Vec, OPA2Vec and the CTD database (ground truth).

**Table 5.** AUC values of ROC curves for chemical–disease association prediction to show the contribution of the ontology-based annotation for literature and ontology descriptions based on the CTD dataset.

|  | Annotated_OPA2Vec | DigChem |
| --- | --- | --- |
| *Accuracy* | 0.873 | 0.841 |
| *Precision* | 0.877 | 0.823 |
| *Recall* | 0.911 | 0.880 |

Chemical–disease associations can typically involve two types of relations: therapeutic, when the chemical is a drug that can be used to treat the disease, and toxic (marker), when the chemical is a toxin causing the disease. To further analyze the contribution of ontology-based text annotation, we extend our experiments to predicting the specific type of the relation (therapeutic or toxic) that associates the chemical and the disease entity by adapting the architecture of our neural network. The results obtained from this experiments are shown in Figure 8 and Table 6. We find that in both types of predictions the normalization based learning, Annotated_OPA2Vec, gives the best performance. In general, we can observe that methods using normalization (Annotated_OPA2Vec, Annotated_Metadata, Annotated_PMC) have better prediction performance than methods which do not use normalization (PMC_only, OPA2Vec).

**Fig. 8:**
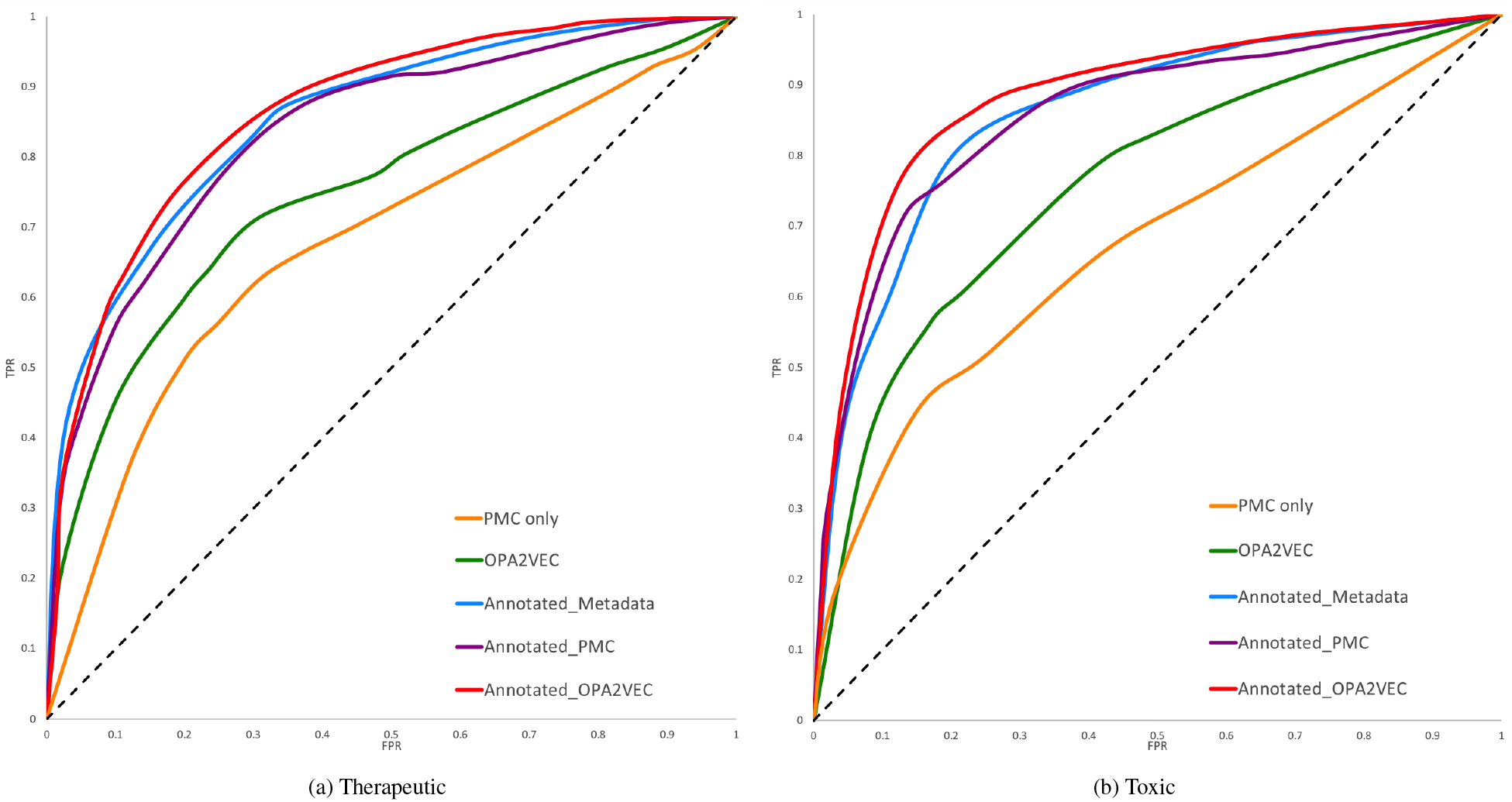
ROC curves for predicting therapeutic and toxic associations between chemicals and diseases in the CTD dataset.

**Table 6.** AUC values of ROC curves for predicting therapeutic and toxic associations between chemicals and diseases in the CTD dataset.

|  | Therapeutic | Toxic |
| --- | --- | --- |
| *PMC_only* | 0.6787 | 0.6689 |
| *OPA2Vec* | 0.7457 | 0.7551 |
| *Annotated_Metadata* | 0.8471 | 0.8519 |
| *Annotated_PMC* | 0.8336 | 0.8531 |
| *Annotated_OPA2Vec* | **0.8594** | **0.8772** |

## 4 Discussion

Ontologies have long been used to provide background knowledge. Recently, this background knowledge is increasingly also incorporated into predictive models through semantic similarity or ontology embedding methods. We have demonstrated that incorporating methods from natural language processing into the feature learning step can further improve the utility of ontologies, mainly through closer alignment between the symbols used in natural language and in formalized theories. While our experiments demonstrate this improvement only in three applications, we believe that it can generalize to other prediction models that rely on combinations of formalized and natural language knowledge.

Our normalization method is currently limited by its reliance on lexical matching to identify mentions of ontology classes, while novel natural language methods often use machine learning models for this purpose as well (Lee *et al.*, 2020). In future work, more experiments with different named entity recognition and normalization approaches are needed to improve our method.

## 5 Conclusion

We have developed a method that integrates information from literature and biomedical ontologies and generates joint embeddings. Using a Siamese neural network, we demonstrated that our method can outperform the-state-of-the-art in several tasks ranging from prediction of interacting proteins through function similarity to prediction of toxicological effects of chemicals. Our experiment results showed that the normalization helps prediction methods to learn previously undetected similarities between biomedical classes and entities which improves the learning and predictive performance.

