## Supplementary for "Self-normalizing learning on biomedical ontologies using a deep Siamese neural network"

*Bioinformatics*
doi.10.1093/bioinformatics/xxxxxx
Advance Access Publication Date: Day Month Year
Manuscript Category

OXFORD

Subject Section

### Supplementary Material:Self-normalizing learning on biomedical ontologies through a deep Siamese neural network

**Fatima Zohra Smaili [1], Xin Gao [1,*] and Robert Hoehndorf [1,*]**

**Abstract**

© The Author 2020. Published by Oxford University Press. All rights reserved. For permissions, please

Table 1. Gene–disease prediction results for orphan diseases where the predicted gene happens to fall within the known disease segment on the right chromosome.

| Disease Name | Disease(OMIM ID) | Gene name | Gene ID | Chromosome | Gene location | Disease segment |
|---|---|---|---|---|---|---|
| Facial Cleft | 119530 | transcription factor AP-2, alpha | MGI:104671 | chr6 | p24.3 | p24.3- |
| Facial Cleft | 119530 | collagen, type XI, alpha 2 | MGI:88447 | chr6 | p21.32 | p24.3- |
| Facial Cleft | 119530 | vascular endothelial growth factor A | MGI:103178 | chr6 | p21.1 | p24.3- |
| Facial Cleft | 119530 | hypocretin (orexin) receptor 2 | MGI:2680765 | chr6 | p12.1 | p24.3- |
| Facial Cleft | 119530 | dystrobrevin binding protein 1 | MGI:2137586 | chr6 | p22.3 | p24.3- |
| Facial Cleft | 119530 | runt related transcription factor 2 | MGI:99829 | chr6 | p21.1 | p24.3- |
| Cleft Palate, Isolated | 119540 | B9 protein domain 1 | MGI:1351471 | chr17 | p11.2 | p11.2- |
| Cleft Palate, Isolated | 119540 | tumor necrosis factor receptor superfamily, member 13b | MGI:1889411 | chr17 | p11.2 | p11.2- |
| Febrile Seizures Familial | 121210 | EYA transcriptional coactivator and phosphatase 1 | MGI:109344 | chr8 | q13.3 | q13-q21 |
| Dyschromatoris Universalis Hereditaria | 127500 | glutamate receptor, metabotropic 1 | MGI:1351338 | chr6 | q24.3 | q24.2-q25.2 |
| Preauricular Fistulae | 128700 | EYA transcriptional coactivator and phosphatase 1 | MGI:109344 | chr8 | q13.3 | q11.1-q13.3 |
| Echo Virus II Sensitivity | 129150 | zinc finger protein 36 | MGI:99180 | chr19 | q13.2 | q13.1-qter |
| Ectrodactyly, Ectodermal Dysplasia and Cleft Lip | 129900 | ankyrin repeat and SOCS box-containing 4 | MGI:1929751 | chr7 | q21.3 | q11.2-q21.3 |
| Ectrodactyly, Ectodermal Dysplasia and Cleft Lip | 129900 | hepatocyte growth factor | MGI:96079 | chr7 | q21.11 | q11.2-q21.3 |
| Immunoglobin A deficiency I | 137100 | runt related transcription factor 2 | MGI:99829 | chr6 | p21.1 | p21.3- |
| Immunoglobin A deficiency I | 137100 | vascular endothelial growth factor A | MGI:103178 | chr6 | p21.1 | p21.3- |
| Immunoglobin A deficiency I | 137100 | hypocretin (orexin) receptor 2 | MGI:2680765 | chr6 | p12.1 | p21.3- |
| Immunoglobin A deficiency I | 137100 | EF-hand domain (C-terminal) containing 1 | MGI:1919127 | chr6 | p12.2 | p21.3- |
| Pseudohypoaldo-steronism | 145260 | calcium channel, voltage-dependent, L type, alpha 1S subunit | MGI:88294 | chr1 | q32.1 | q31-q42 |
| Malignant Hyperthermia | 154275 | keratin 16 | MGI:96690 | chr17 | q21.2 | q11.2-q24 |
| Malignant Hyperthermia | 154275 | hypocretin | MGI:1202306 | chr17 | q21.2 | q11.2-q24 |
| Malignant Hyperthermia | 154275 | neurofibromin 1 | MGI:97306 | chr17 | q11.2 | q11.2-q24 |
| Malignant Hyperthermia | 154275 | erb-b2 receptor tyrosine kinase 2 | MGI:95410 | chr17 | q12 | q11.2-q24 |
| Malignant Hyperthermia | 154275 | signal transducer and activator of transcription 3 | MGI:103038 | chr17 | q21.2 | q11.2-q24 |
